# How does a small molecule bind at a cryptic binding site?

**DOI:** 10.1101/2021.03.31.437917

**Authors:** Yibing Shan, Venkatesh P. Mysore, Abba E. Leffler, Eric T. Kim, Shiori Sagawa, David E. Shaw

## Abstract

Protein-protein interactions (PPIs) are ubiquitous biomolecular processes that are central to virtually all aspects of cellular function. Identifying small molecules that modulate specific disease-related PPIs is a strategy with enormous promise for drug discovery. The design of drugs to disrupt PPIs is challenging, however, because many potential drug-binding sites at PPI interfaces are “cryptic”: When unoccupied by a ligand, cryptic sites are often flat and featureless, and thus not readily recognizable in crystal structures, with the geometric and chemical characteristics of typical small-molecule binding sites only emerging upon ligand binding. The rational design of small molecules to inhibit specific PPIs would benefit from a better understanding of how such molecules bind at PPI interfaces. To this end, we have conducted unbiased, all-atom MD simulations of the binding of four small-molecule inhibitors (SP4206 and three SP4206 analogs) to interleukin 2 (IL2)—which performs its function by forming a PPI with its receptor—without incorporating any prior structural information about the ligands’ binding. In multiple binding events, a small molecule settled into a stable binding pose at the PPI interface of IL2, resulting in a protein–small-molecule binding site and pose virtually identical to that observed in an existing crystal structure of the IL2-SP4206 complex. Binding of the small molecule stabilized the IL2 binding groove, which when the small molecule was not bound emerged only transiently and incompletely. Moreover, free energy perturbation (FEP) calculations successfully distinguished between the native and non-native IL2–small-molecule binding poses found in the simulations, suggesting that binding simulations in combination with FEP may provide an effective tool for identifying cryptic binding sites and determining the binding poses of small molecules designed to disrupt PPI interfaces by binding to such sites.

## Introduction

Protein-protein interactions (PPIs) play an important role in mediating many biological processes, and the dysregulation of such interactions is central to the development of numerous human diseases.^1^ Targeting disease-associated PPIs with highly selective small drug molecules is a promising therapeutic strategy.^2^ Developing drugs that target PPIs, however, has proven highly challenging.^3–5^ Whereas conventional drugs often bind to well-defined pockets or clefts that are present even in the absence of a ligand, binding sites at PPI interfaces tend to be “cryptic”: In the absence of a ligand, cryptic binding sites predominantly adopt a relatively flat and featureless conformation, but when bound to a ligand, they resemble traditional binding pockets.^6^ Because cryptic pockets are not readily visible in the unbound protein, the ability to accurately predict the locations, shapes, and chemical characteristics of these pockets through computational studies could provide a highly advantageous starting point for the structure-based design of PPI inhibitors.

Molecular dynamics (MD) simulation has become a promising tool for the identification and characterization of cryptic binding sites.^7–10^ An increasing number of studies in recent years have set the goal of using MD to characterize the conformational dynamics of known cryptic ligand-binding sites starting from known bound structures, or to identify their locations from apo structures.^11–19^ It is yet to be demonstrated, however, that MD simulations are capable of recapitulating the binding process of a small molecule to its correct pose in a cryptic site without incorporating experimental information about the bound structure. We have previously reported unbiased MD simulations of the complete binding processes of several non-cryptic smallmolecule drugs to their respective kinase targets;^20,21^ these simulations successfully found the native binding poses without any input based on structural knowledge of the bound state. To achieve a similarly detailed unbiased characterization of cryptic binding is more challenging, however, because it requires accurate simulation of both ligand binding and the formation of the binding site itself.

Here, we report the results of unbiased, all-atom MD simulations of cryptic-site binding of the small molecule SP4206 and three of its analogs with interleukin 2 (IL2). IL2 is a cytokine that interacts with the IL2 receptor (IL2R) to regulate the activity of white blood cells in the immune response,^22,23^ and SP4206 is a candidate drug molecule that inhibits IL2 activity by binding to IL2 (dissociation constant K_d_ = 60 nM^24^) in a cryptic binding site at its interface with the α unit of IL2R.^25^ In our simulations, we observed the full process of IL2 binding to SP4206 and to three SP406 analogs, with each small molecule reaching a binding site and pose virtually identical to those in the IL2-SP4206 crystal structure without the use of any prior structural information about their binding. The small molecule in each simulation was placed at a random initial position distant from IL2, and arrived at the native binding pose (Movie S1) as a binding groove simultaneously emerged at the cryptic binding site, resulting in a stable protein-ligand complex that is virtually identical (Figure 1) to the crystal structure of the complex.^26^

**Figure 1.**
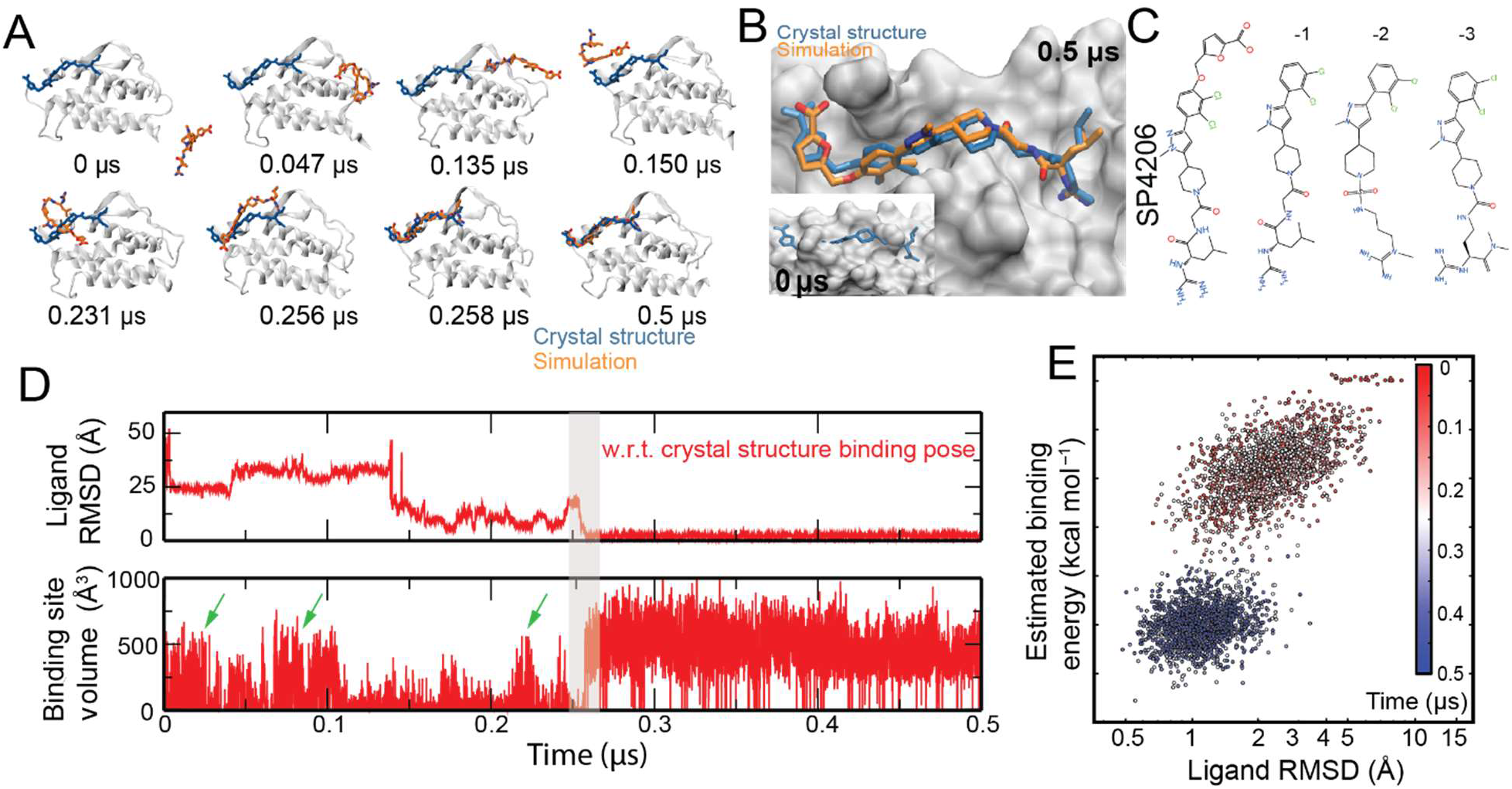
Simulation of SP4206 binding to IL2. (A) Snapshots taken from Simulation 1, in which one of the three SP4206 ligands in the system reached the native binding pose with IL2. The crystal pose (PDB 1PY2) of the ligand (blue) is also shown for reference. (B) The simulation ligand-binding pose compared with the crystal pose. Also shown are conformations of the protein before (0 μs, inset) and after (0.5 μs) binding, in surface representation. Notably, the binding groove is not present at the initial conformation of the simulation. (C) Chemical structures of SP4206 and analogs. (D) Time series of the SP4206 RMSD with respect to the crystal binding pose in juxtaposition with the series of the volume of the binding groove. Note that at the SP4206-binding site, a transient groove that is comparable in size to the native binding groove emerged at approximately 0.02 μs, 0.08 μs, and 0.22 μs (marked by green arrows), prior to the ligand binding at 0.25 μs (gray area). (E) The binding process described by estimated binding energy (*y*-axis) and conformational fluctuation of the ligand, as measured by RMSD with respect to the conformation of the previous time step (*x*-axis). Additional analysis of the simulations is presented in Figure S2A.

When a small molecule binds to a protein, the ligand and protein first form encounter complexes, in which the ligand is located in non-native binding sites or in the native binding site but in non-native binding poses. Encounter complexes often last tens of microseconds or longer, beyond the timescale of the simulations we performed. To distinguish the native protein–small-molecule complex structure from the non-native encounter-complex structures, we analyzed the structures in our simulations using free energy perturbation (FEP)^27^ calculations to estimate the binding free energy. We found that these FEP calculations of the binding free energy were sufficiently accurate to distinguish the native binding pose from the encounter complexes, with the native pose consistently having greater binding free energy. Taken together, our results suggest that unbiased MD simulations combined with FEP may provide a useful means of predicting the locations of cryptic binding sites and the binding poses of small molecules in these sites.

## Results

### Unbiased simulations of ligand binding at a cryptic site

We started by performing six simulations of IL2 binding SP4206 (a total of 134 μs of simulation time, with each simulation containing either one or three copies of SP4206; see Table S1 for details). SP4206 bound IL2 in the native binding site and remained stable in the native binding pose in two of these simulations. (Transient non-native interaction of SP4206 with the native binding site was also observed in the simulations.) In one simulation (Simulation 1, a 31-μs simulation with three copies of SP4206), SP4206 bound in the native binding pose after 0.25 μs of simulation time (Figure 1D and Movie S1). In another simulation (Simulation 3, an 18-μs simulation with one copy of SP4206), binding occurred after 0.1 μs. In both of these simulations, SP4206 diffused extensively in the space around IL2 before settling into its binding site (Figure 1A). The binding poses generated in both of these simulations are virtually identical to the native binding pose captured in the crystal structure (Figure 1B), with an average ligand root-mean-square deviation (RMSD) of 1 Å from the crystal structure pose (Figure 1E). After reaching the native binding pose, SP4206 remained highly stable (Figure 1D) for the remainder of these two simulations. At the beginning of the simulations, the cryptic binding site was largely flat. A stable binding groove only emerged during the binding process (Figure 1B and 1D). Conformational fluctuation of the cryptic binding site was greatly reduced after SP4206 binding (Figure S1). The emergence of the binding groove is discussed in greater detail below.

To better gauge how frequently native binding occurs in simulations, we then launched 10 additional 10-μs unbiased simulations (Simulations 7–16; see Table S1) of SP4206 and IL2, each with one copy of the small molecule and one copy of the protein. In four of the 10 simulations, SP4206 reached the native binding pose, after 1.05, 0.4, 0.6, and 8.0 μs of simulation time (Figure 1F).

We then performed unbiased simulations (Simulation 17–24) of three analogs of SP4206 (SP4206-1, SP4206-2, and SP4206-3)^28^ with IL2. Because these analogs bind IL2 more weakly (K_d_ = 2, 4, and 7 μM, respectively), we included two copies of the small molecule in each simulation to increase the concentration. Despite the weaker binding, each analog reached and remained stable at the native binding pose in multiple simulations (Figure 2A). SP4206-1 (Simulations 17 and 18) reached the native binding pose in two out of two 20-μs simulations (at 16 and 0.1 μs, respectively), SP4206-2 (Simulations 19 and 20) in two out of two 20-μs simulations (at 1 and 0.6 μs, respectively), and SP4206-3 (Simulations 21–24) in two out of four 20-μs simulations (Simulation 22 at 15 μs and Simulation 24 at 6.1 μs). The binding processes of the analogs were similar to those observed in the SP4206 simulations, including the opening of the binding groove and rearrangement of the side chains at the binding site.

**Figure 2.**
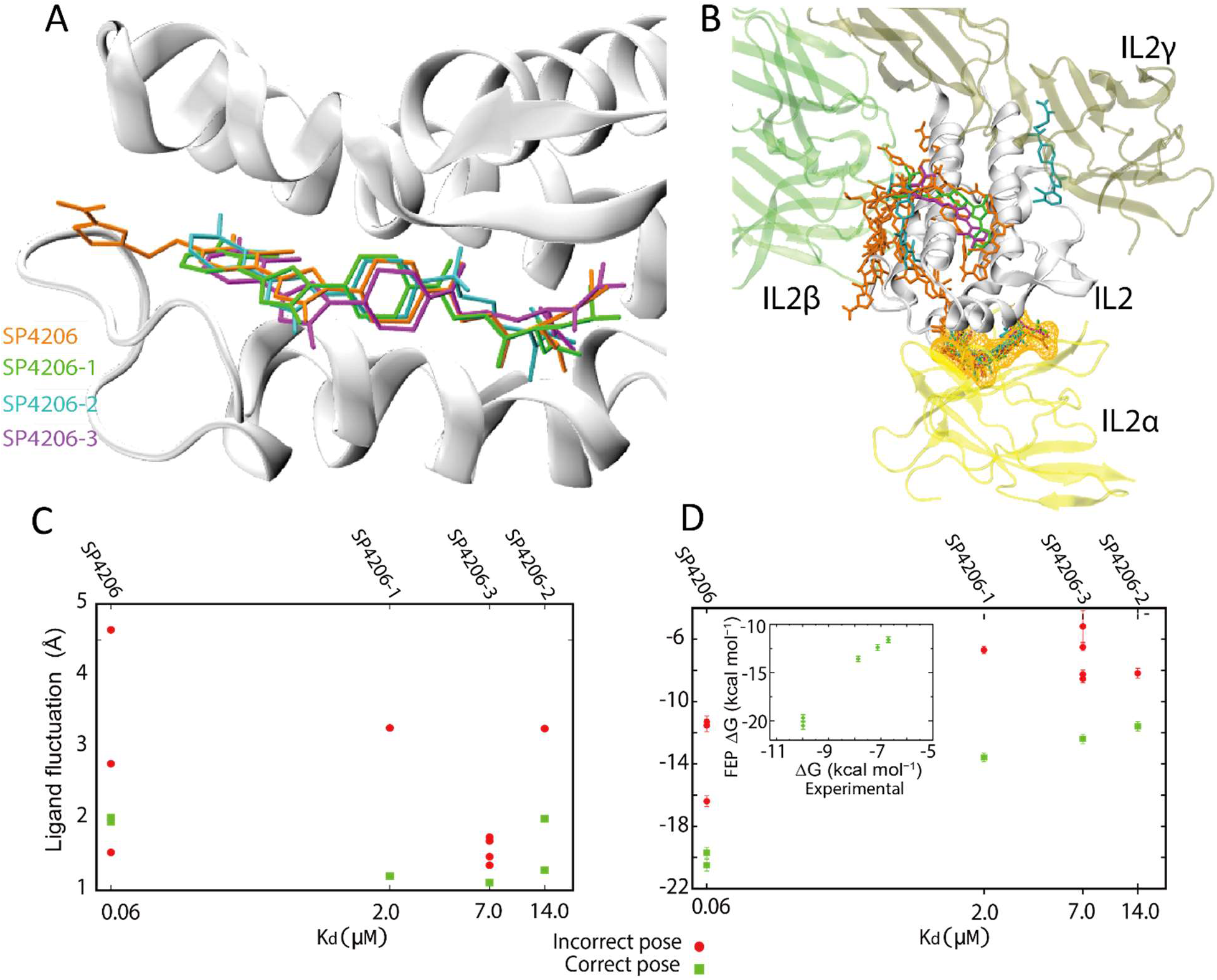
Simulations of binding of SP4206 analog molecules and free energy calculation. (A) The binding poses of SP4206 and its analogs generated by spontaneous binding simulations. (B) The non-native binding poses of SP4206 and its analogs generated by simulations, shown in the structural context of IL2-IL2R association. The native binding pose is also shown with a mesh around the small molecule. The colors of the small molecules are as shown in (A). (C) The conformational fluctuations of SP4206 and analogs in simulation-generated binding poses (*y*-axis) compared to the dissociation constants of the compounds (*x*-axis). As shown, with one exception for SP4206, the fluctuation is smaller in the native binding pose than in the non-native ones for the same compound. (D) The FEP binding free energies of SP4206 and analogs in various simulation-generated binding poses (*y*-axis) compared to the dissociation constants (K_d_) of the compounds (*x*-axis). The binding free energy was estimated to be 20.5 ± 0.38 or 19.7 ± 0.33 kcal mol^-1^ for SP4206, 13.58 ± 0.28 kcal mol^-1^ for SP4206-1, 11.59 ± 0.29 or 11.58 ± 0.28 kcal mol^-1^ for SP4206-2, and 12.39 ± 0.3 kcal mol^-1^ for SP4206-3. In the inset, the calculated binding energy (*y*-axis) is compared to the binding energies derived from K_d_ (*x*-axis). For a given compound, the calculated binding energy is consistently greater for the native binding pose than for the non-native ones.

### Distinguishing the native binding poses from the non-native ones

When bound to its heterotrimeric receptor IL2R (K_d_ ≈ 10 pM), IL2 interacts with the receptor’s α, β, and γ chains at three distinct PPI interfaces.^29^ SP4206 inhibits IL2 activity by binding at the α interface and disrupting IL2’s interactions with IL2R. Based on crystallographic structures and other experimental evidence, the native binding of SP4206 and its analogs at the α interface is known to be highly specific, and this was also observed in our simulations. In addition to the native binding, SP4206 and its analogs did become trapped in non-native poses at or near the β and γ interfaces in some of our simulations (Figure 2B), most likely because the characteristics of PPI interfaces (e.g., the exposed hydrophobic surface and the prevalence of bridging water molecules) make them prone to interactions with small molecules, including nonspecific interactions.^4^

Summing up the total time before a binding event was observed across all of our simulations, and factoring in the small-molecule concentration in each simulation, our simulations suggest a binding on-rate on the order of 4 μM^-1^s^-1^. For SP4206, with its K_d_ of 60 nM, the off-rate would thus be ~0.24 s^-1^, or on average ~4.2 s per dissociation event. It would thus be completely impractical to directly simulate the dissociation of SP4206 or its analog inhibitors from IL2 in order to obtain the dissociation constant. Even some non-native states dissociate too slowly for the dissociation time to be measured in simulation: For each of the four small molecules we simulated, we observed at least one instance of non-native binding in which the molecule was trapped for at least 5 μs, and in none of these instances did the molecule subsequently dissociate. Such slow dissociation is to be expected, as even assuming a fast diffusion-limited on-rate for non-native binding of ~100 μM^-1^s^-1^,^30^ and a relatively weak dissociation constant of ~100 μM, the residence time would be ~100 μs. It is thus impractical to directly measure dissociation time in simulation as a way to distinguish native from non-native binding.

From a practical point of view, however, the ability to distinguish native from non-native binding is crucial for the application of unbiased MD simulations of small-molecule binding to drug discovery. FEP is a theoretically rigorous calculation that explicitly takes into account, among other factors, the effect of water molecules in both small-molecule solvation and binding. Its application, however, has been hampered by force field inaccuracies and by incomplete sampling due to limitations in computational resources. With the continuing growth of computational power, the development of new sampling methodology, and improvements in force field accuracy, the accuracy of FEP calculation of ligand-binding free energy has improved substantially in recent years.^31^ We calculated using FEP^32^ the binding free energies of various stable IL2 binding poses adopted by small molecules in our simulations. Because for a potent small-molecule binder like SP4026 the binding free energy of non-native binding is substantially less than that of native binding, we reasoned that FEP should be able to identify the native binding pose despite the fact that the accuracy of FEP calculations may vary.

We found that distinguishing native from non-native binding for the protein–small-molecule system examined here appears to be feasible, despite the known difficulties of converging binding FEP calculations. Our FEP calculations showed that the binding free energy associated with a non-native binding event is consistently lower than the binding free energy of native binding events of the same small molecule (Figure 2D). To make the comparison, we first performed an FEP calculation for each native binding pose observed in our simulations. Two native binding events each were observed for SP4206 and SP4206-2, and—reassuringly—we found that the binding free energies estimated from the two independent binding events were highly consistent. The calculated binding free energies of the four small molecules correlate well with their K_d_ values. The experimental binding free energies derived from the K_d_ (ΔG = –*RT*ln[K_d_], with R denoting the gas constant and T temperature) are considerably weaker than, but well correlated with, the FEP estimates (inset of Figure 2D). Importantly, the nonnative binding free energy is significantly lower than that of native binding (Figure 2C), showing that FEP correctly distinguished native and non-native binding poses.

Because the computational cost of these FEP calculations is relatively high, we decided to also test the less rigorous, but less computationally intensive Generalized Born (GB) model augmented with the hydrophobic solvent accessible surface area (SA) term (the GBSA model)^33^ to estimate the binding free energy. (In the GBSA method, unlike in FEP, water molecules are not represented explicitly, and their effects are accounted for using an implicit model.) In applying the GBSA calculation to snapshots generated by our simulations, we found that the GBSA binding energy is generally higher for native than for non-native binding, but not always. Non-native binding in which the small molecule is highly buried in IL2 can lead to GBSA estimates for the binding energy that are higher than for native binding. To improve the ability of the GBSA method to distinguish the native and non-native binding, we then examined the overall pattern of the energy distribution in binding. Reminiscent of the two-state underlying energy landscape often seen in protein folding,^34^ we found that the native binding process tended to feature a clear bimodal distribution of the GBSA energy and a sharp rise of the binding energy upon adoption of the native binding pose (Figures 1E and S2D), which is consistent with cooperative formation of the native binding contacts. We found that, in contrast to the native binding process, the GBSA binding energy for non-native binding tended to follow unimodal patterns (Figure S2), suggesting that cooperativity was absent for the non-native interactions. Moreover, with the exception of one non-native binding event, we found that SP4026 exhibited less conformational fluctuation in native than in non-native binding in our simulations (Figure 2D).

### The binding pathway and the putative transition state

In some protein-protein binding processes, electrostatic interactions play an important role in guiding protein diffusion and speeding up formation of the native complex.^35^ Such interactions were important in the binding of IL2 and SP4206 in our simulations; SP4206 carries an electric dipole with a negatively charged furonic acid group at one end and a positively charged guanido group at the other. In our simulations, we observed that, prior to binding, SP4206 lingered in the vicinity of the native binding site, forming encounter complexes with IL2. In these encounter complexes, the orientation of the small molecule was fairly heterogeneous, but with a bias toward a native-like orientation that appears to be due to electrostatic interactions with IL2 (Figure 3A). This suggests that electrostatic interactions with IL2 speed up the binding of SP4206 in the native pose.

**Figure 3.**
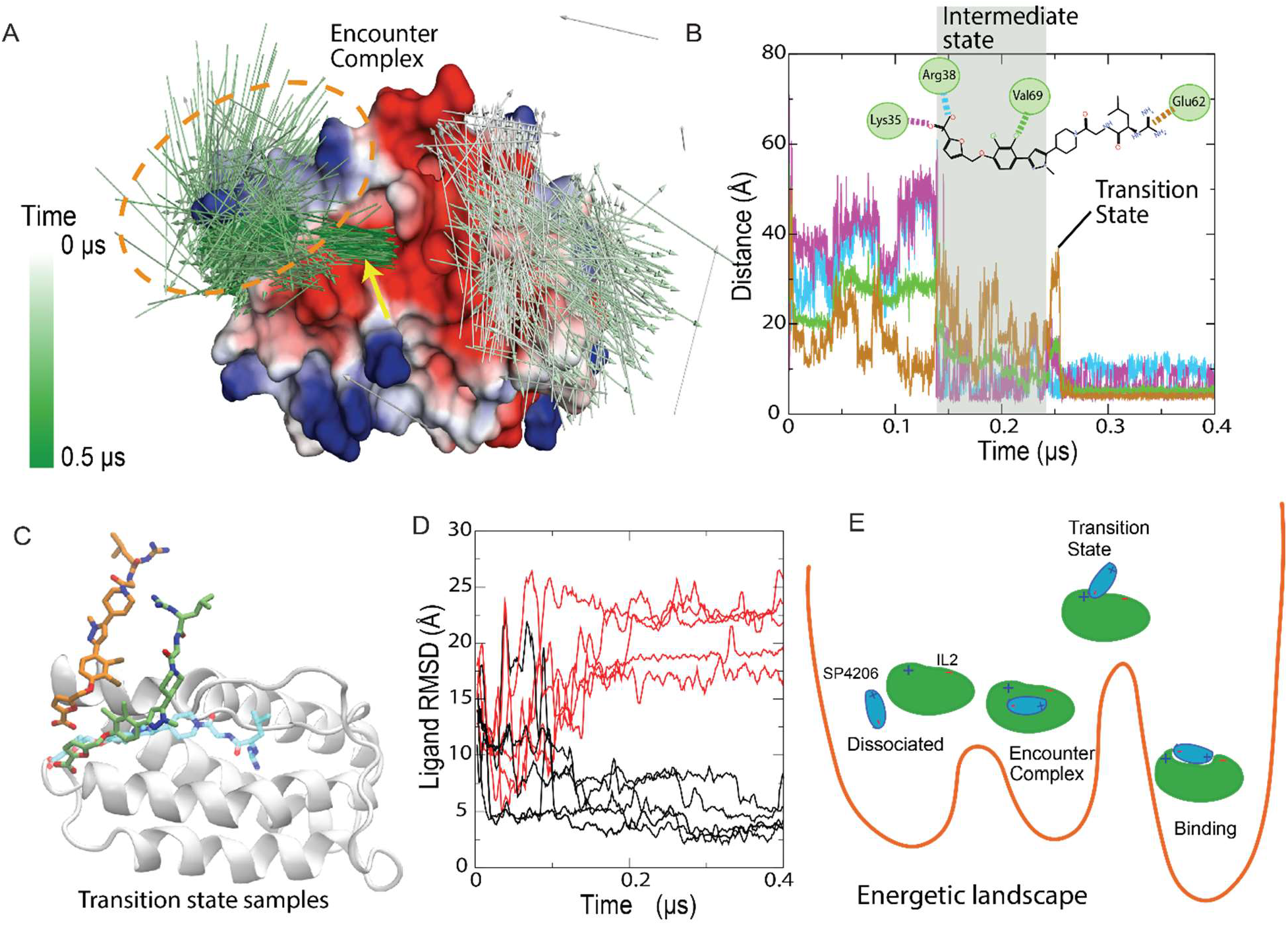
Binding pathway and the transition state. (A) The binding process shown in Figure 1 in terms of SP4206 dipole directions, which are color coded according to simulation time. In the intermediate state (left cluster, circled in orange) the arrowheads tend to be pointing to the left, reflecting a general alignment of the dipole directions; the well-aligned arrowheads (indicated by the yellow arrow) just underneath this cluster show the native binding pose of SP4206. The surface of IL2 is colored by the local electrostatic properties of IL2. (B) Evolution of IL2-SP4206 contacts (defined in the inset) in the binding process. (C) Two conformations identified from binding simulations as members of the transition state ensemble. The native pose is also shown. (D) 10 simulations launched from the orange conformation in (C), of which 5 (black lines) quickly led to native binding. (E) A sketch of the energetic landscape and pathway of the binding.

The crystal structure of the IL2-SP4206 complex suggests that three key interactions between the two molecules contribute to the binding free energy: the ion pairs of the furonic acid with Lys35 or Arg38, the burial of the dichlorobenzene, and the ion pair between the guanido group and Glu62 (inset of Figure 3B). In our simulations, these three interactions by and large formed sequentially, from one end of SP4206 to the other: The interaction with the furonic group often formed first, serving as an anchor and restraining the small molecule to the vicinity of the native binding site. Subsequently, the dichlorobenzene group approached its native binding position, while the guanido group was still distant from its native position (Figure 3B). The guanido group eventually fell into its native interaction with Glu62, marking the completion of the binding event (Figure 3B). In the simulations, we also occasionally observed a slightly different process of SP4206 binding, in which the guanido interaction formed first, and the furonic acid interaction last. In all of the binding events observed in the simulations that led to native binding, however, the first protein–small-molecule native interaction that formed was electrostatic in nature.

The simulations suggest that the loss of some native SP4206-IL2 contacts in the encounter complex, while maintaining a native ion pair, may be characteristic of the rate-limiting step of binding. We conjectured that these conformations, in which SP4206 is largely (with the exception of the ion pair) not in contact with IL2, belonged to the transition state ensemble (the state in the binding pathway from which there is a 50% probability of reaching the end state) (Figure 3C). These conformations are reminiscent of those seen in transition states in proteinprotein binding, where some native contacts are formed, but the interfaces are mostly exposed to solvent.^36^ The solvent exposure of SP4206 in this state is reflected in the fact that, immediately prior to binding, its RMSD from the final binding pose increased significantly (Figures 1D and 3B)

To test whether the transition state ensemble does indeed contain conformations like this (Figure 3C), we picked one such conformation from our binding simulations and from it started a set of 10 10-μs simulations. In five of these simulations, SP4206 arrived at or near the native binding pose within 200 ns, and in the other five simulations SP4206 drifted away from the native biding pose within 200 ns and did not reach the native binding pose by the end of the simulation (Figure 3D). While this exact 50% binding probability is fortuitous, the result supports the notion that the initial conformation is indeed part of the transition state ensemble. A more comprehensive characterization of transition state conformations is, however, beyond the scope of this study, and will require a dedicated investigation with a much greater number of simulations that start from a more diverse set of conformations.

### The emergence of the binding groove

It has long been debated whether the emergence of a binding pocket or groove in a cryptic binding site is a process of induced fit or one of conformational selection.^37^ In an induced-fit process, the binding site adopts the bound conformation only upon interaction with the ligand. In a conformational-selection process, on the other hand, the binding site transiently adopts its bound conformation in a dynamic equilibrium with other conformations—independent of interactions with the ligand—and ligand binding is conditional upon the adoption of the bound conformation by the host protein.^38^ Many binding processes, however, may have a hybrid character, with some native ligand-protein interactions first forming as an “anchor” (as in conformational selection), and “latch” interactions subsequently forming in ways consistent with an induced-fit process.^39^

In our simulations, a groove intermittently emerged at the binding site even without a small molecule nearby, and the groove occasionally reached volumes comparable to those of the SP4206-binding groove observed in the bound state (e.g., at 25 ns and at 100 ns in Simulation 1; Figure 1D and Movie S1). This spontaneous formation of the groove presumably resulted from the clustering of solvent-exposed hydrophobic residues (e.g., Phe42, Phe44, Tyr45, and Leu72) at the binding site and the relatively poor local packing. Without an occupying ligand, however, the groove was transient and highly variable in shape (Figures 4A and S1). As noted in the discussion of the putative transition state, in the IL2-SP4206 binding process a key initial step was often the formation of the ion pair between the furonic group of SP4206 and Arg38. The arginine in our simulations of apo IL2 alternated between two conformers (Figure 4C): In one, the side chain was adjacent to Phe42, and in the other the side chain was adjacent to Lys35. In our simulations, this ion pair preferentially formed when IL2 was in the latter conformer, because in this conformer Arg38 was more exposed and the local positive electrostatic potential was reinforced by Lys35.

**Figure 4.**
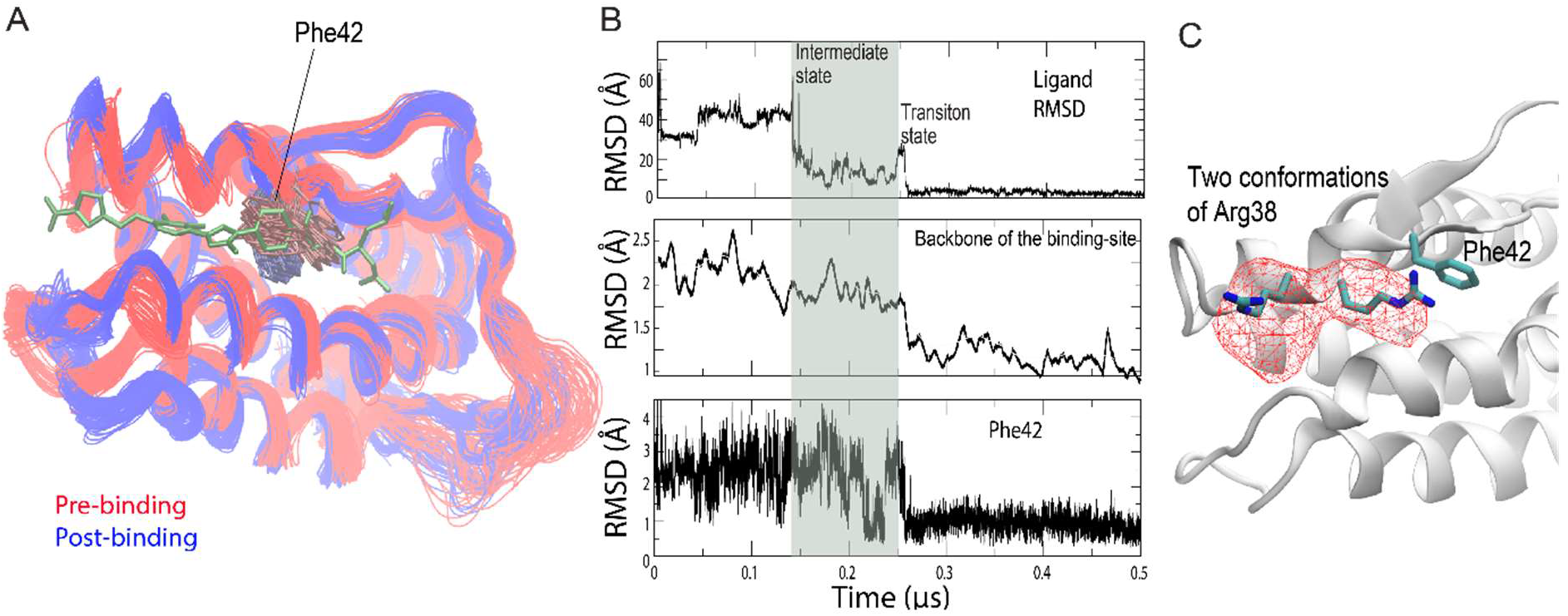
Emergence of the binding groove. (A) Conformations of the IL2 backbone and Phe42 before and after SP4206 binding. Multiple conformations from the binding simulation discussed in Figures 1 and 3 are superimposed. The backbone of the residues lining the long-side edges of the SP4206 binding groove moved further apart, and Phe42 was stabilized in a single rotamer after binding. (B) Top panel: the time series of SP4206 RMSD with respect to the crystal binding pose (identical to the top panel of Figure 1D); middle panel: two running averages (of different widths of the averaging window) of the RMSD of the backbone of the residues lining the long-side edges of the binding groove (residues 32–44, 63–74) with respect to the SP4206-bound conformation; bottom panel: RMSD of Phe42 with respect to the SP4206-bound conformation. (C) The spatial occupancy (mesh) of the two conformers of Arg38 from a 5-μs simulation of apo IL2, in which SP4206 and other small molecules were not present.

The formation of the initial ion pair between SP4206 and IL2 can thus be seen as characteristic of a conformational-selection process, since this interaction preferentially formed when Arg38 adopted the latter conformer. The rest of the process, however, appears to be better characterized as induced fit. As shown in Figure 4A, upon SP4206 binding the binding groove widened: the backbones of the residues lining the two opposite sides of the binding groove were farther apart than before binding, and the side chains of these residues (notably Phe42) assumed different conformations (Figure S3). The residues approached their native binding conformation only after the small molecule arrived at the vicinity of the binding site. As shown in Figure 3B (from 0.14 μs to 0.26 μs), SP4206 maintains extensive, albeit unstable contacts with the binding site. The presence of SP4206 reduces the conformational fluctuation of the binding site, and it was only after SP4206 adopted the native pose that Phe42 and the backbones of the binding-site residues fully settled at the native binding conformation (Figure 4B). This order of events is consistent with the notion that non-native interactions with the small molecule lower the energetic barrier for the conformational change.^7^ The nanosecond-timescale separation between the two events is consistent with the fact that side-chain rearrangements commonly occur on that timescale.^40^ From our simulation observations, we concluded that the SP4206 binding of IL2 is a hybrid process of both conformational selection and induced fit, which explains why a stable binding groove at the cryptic site is not observed in simulations in the absence of the ligand.

### IL2 interactions with chemical fragments of SP4206

SP4206 was originally generated by the assembly and optimization of two weakly binding chemical fragments first identified by site-specific fragment screening against IL2.^24^ Such fragment-based approaches show great promise for generating high-affinity lead compounds. ^41^ To investigate how the interactions involved in the IL2 binding of SP4206 are inherited from the IL2 interactions of the chemical fragments, we broke down SP4206 into chemical fragments and simulated their spontaneous binding. We first broke SP4206 into two fragments (S and T) of approximately equal size (Figure 5A) and performed unbiased simulations of these two relatively large fragments with IL2 (Movie S2). In the simulations (Simulation 25 (24 μs) and 26 (53.9 μs)), these relatively large fragments retained their binding specificity: Each reached binding locations consistent with, and binding poses approximately consistent with, those of SP4206 (Figure 5A), although they did not remain there stably to the end of the simulations. In Simulation 26, for instance, fragment T remained in the native binding pose for ~25 μs, but dissociated from IL2 before the end of the simulation.

**Figure 5.**
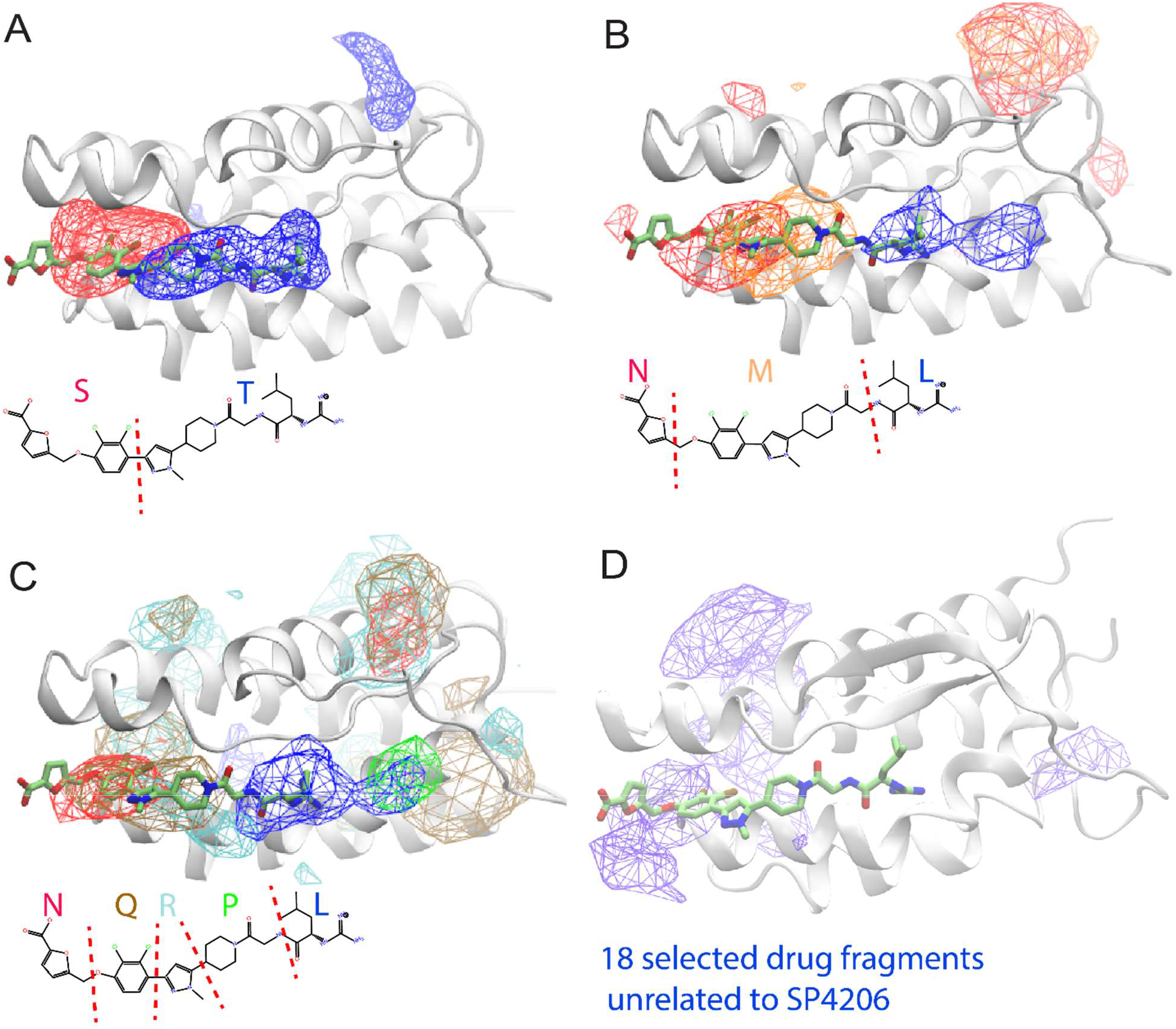
Binding of fragments of SP4206 to IL2. The occupancy density maps of fragments of SP4206: S and T in (A) based on a 54-μs simulation; N, M, L in (B) based on a 37-μs simulation; and N, Q, R, P, L in (C) based on a 16-μs simulation. In (D), the occupancy density of 18 common drug fragments unrelated to SP4206 is shown, based on a 16.5-μs simulation. The native binding pose of SP4206 is shown, to mark the cryptic binding site and to indicate the locations of the fragments relative to a bound SP4206 molecule.

We then broke SP4206 into three fragments (N, M, and L). In an unbiased simulation with IL2 (Simulation 27 (37 μs), Movie S3), the three fragments also reached the approximate binding site, but they did not settle into any well-defined binding poses. This was particularly the case for the N and L fragments, which were smaller than fragment M (Figure 5B).

Finally, we broke SP4206 into five very small fragments (N, Q, R, P, and L) and performed an unbiased simulation (Simulation 28 (16.2 μs)) of them with IL2 (Movie S4). Although these small fragments almost completely lost binding specificity with IL2, the five fragments noticeably clustered near the location of the SP4206 binding site, outlining it (Figure 5C). As with SP4206, the fragments also visited locations other than the cryptic binding site, with the fragments (due to their lower binding specificity) spending more time away from the site than did the full SP4206 molecule. The interaction of the fragments with the cryptic binding site was transient: None of the fragments remained in the binding site for more than a few μs, reflecting the much weaker binding affinity they had for IL2 than did SP4206. However, even for the five smallest fragments, statistical analysis indicates that the cryptic binding site remained the most commonly visited location (Figure 5C).

These simulations of fragment binding suggest that, although fragments taken from a high-affinity small molecule gradually lose the specificity in their interactions with the host protein as they decrease in size,^42^ they still preferentially interact with the binding site, presumably recognizing the electrostatic properties, the exposed hydrophobic surface, and the relatively poor local packing. We further simulated IL2 with (Simulation 29 (16.5 μs)) 18 fragments (of comparable size to the SP4206 fragments, with molecular weights ranging from 104 to 397) identified as common in oral drug molecules,^43^ and found that they clustered at the cryptic binding site to a lesser degree than did the SP4206 fragments (Movie S5 and Figure 5D). These observations confirm the notion that identifying chemical fragments that interact favorably with a cryptic binding site may help narrow down the chemical space in which to search for cryptic-site inhibitors in drug discovery. In the meantime, it appears that attempts to design cryptic small-molecule inhibitors by assembling fragments observed to bind in simulations are likely to work best when the chemical fragments are of sufficient size to exhibit specific and transferable native interactions with proteins.

### Simulation of another case of ligand binding at a cryptic site

Our work raises the question of whether unbiased MD simulations of protein–small-molecule binding are likely to be generally applicable to other protein-ligand systems, or if IL2-SP4206 is in some way unusually well suited to this approach. SP4206 carries, for example, a large dipole moment and forms two ion pairs with IL2 in binding. It is thus possible that the success of unbiased MD simulations of IL2-SP4206 binding are a consequence of the prominent role of the electrostatic interactions in the binding. To test whether this is the case, we decided to simulate a system known to feature cryptic-site binding that does not have such strong electrostatic interactions. Compound 43a (Figure 6B) is a sub-μM lead molecule^44^ that precedes ABT-199, an approved drug molecule for treatment of chronic lymphocytic leukemia. Binding of compound 43a to Bcl-2 and Bcl-xL at a cryptic binding site disrupts their protein-protein interfaces. In two of our six unbiased simulations (a total of 148.5 μs of simulation time) of 43a binding to Bcl-xL (see Methods), two native binding events were observed (Figure 6), which generated binding poses that strongly resemble the crystal structure (PDB 2O2M). Although, as in SP4206 binding, significant conformational changes were observed at the cryptic binding site during the binding process of 43a, in contrast to SP4206 binding electrostatic interactions did not appear to play a crucial role. Although definitive answers would require further investigation on more cases of protein–small-molecule binding, these results suggest that unbiased MD simulation may be broadly applicable in the identification of cryptic sites and the characterization of cryptic-site binding.

**Figure 6.**
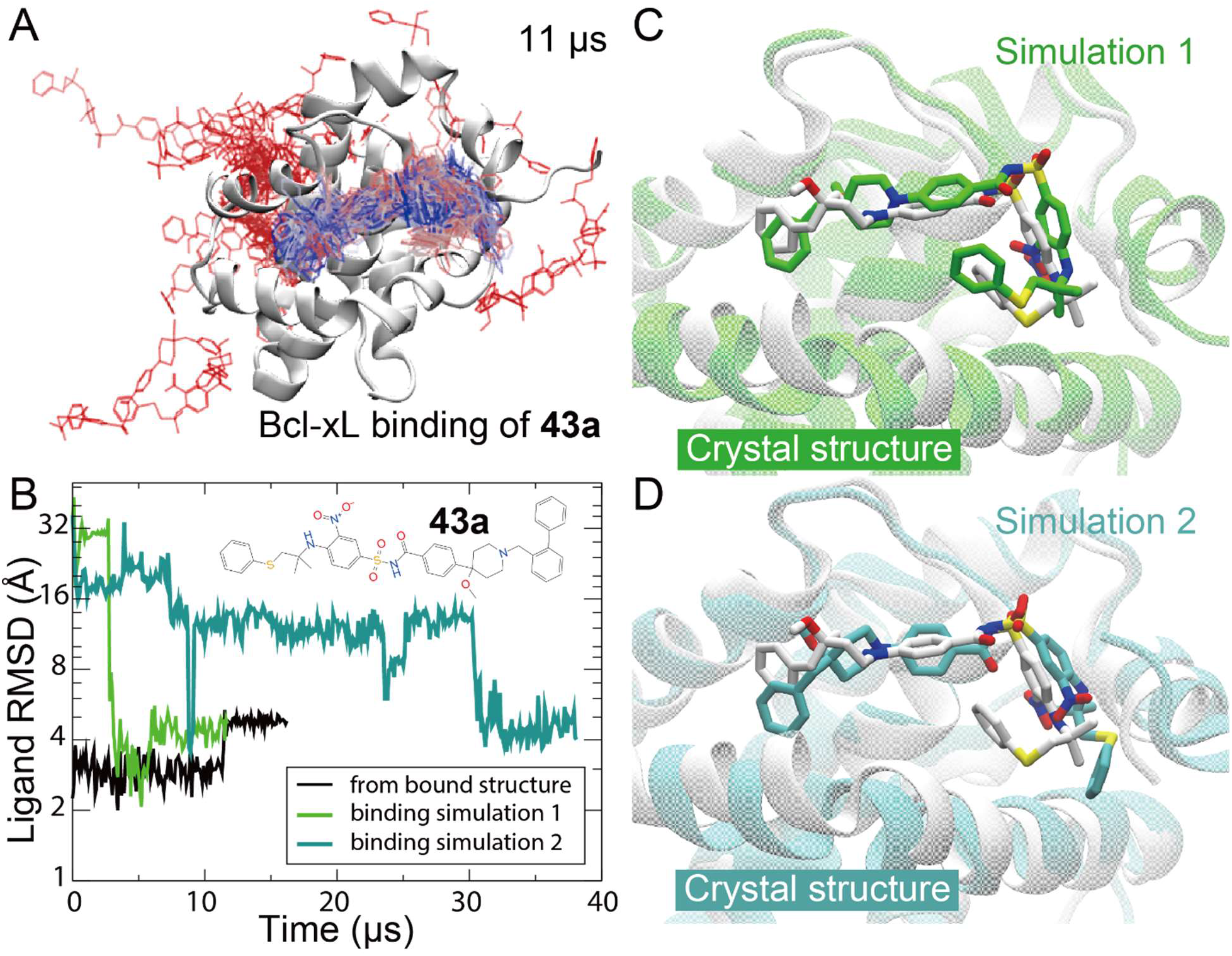
Compound 43a binding to Bcl-xL. (A) Positions of the small molecule in an 11-μs simulation (Simulation 1) in which it reached the native binding pose. Simulation time is color-coded from red, to gray, to blue. (B) The RMSD of the small molecule with respect to the crystal structure (PDB 2O2M) in the binding processes of Simulations 1 (green) and 2 (cyan). Also shown is the RMSD of the small molecule in a simulation starting from the bound structure. All three simulations eventually settled at the same RMSD region. (C) and (D) The binding poses generated by Simulations 1 and 2, respectively, compared with the crystal structure. Note that simulation-generated binding poses are very similar but not identical to the crystal structure. The relatively minor discrepancies may be attributable to the force field for the small molecule.

## Discussion

One of the major challenges of targeting cryptic binding sites in drug discovery is identifying the location of the binding site and determining the early lead molecule’s binding pose. In the absence of such structural information, attempts to improve binding affinity and other properties with medicinal chemistry can be inefficient, and often fruitless. The relatively low binding affinity of pre-optimization lead molecules can make it difficult to attain structural information by conventional structural biology methods such as X-ray crystallography. The work reported here suggests that, given the rapid growth in computational power and steady improvement in simulation force fields, the combination of simulations of spontaneous small-molecule binding together with FEP calculations may serve as an effective computational platform for the structural characterization of protein–small-molecule interactions in the early stages of drug discovery projects seeking cryptic-site inhibitors. In concert with nuclear magnetic resonance (NMR) or surface plasmon resonance (SPR) screening, spontaneous small-molecule binding simulations may help reveal the locations of cryptic sites and the poses of binders therein, and FEP calculations can be used to distinguish the non-native binding events observed in the simulations. This computational approach may be especially effective in providing insight into ways to combine small molecules into large and more potent inhibitors in conjunction with the widely used “structure-activity relationships (SAR) by NMR” method.^45^

Although SP4206 did not bind at the native binding site in all of our simulations, it never settled in the native site in a non-native pose—a finding that illustrates the highly specific nature of the native IL2 binding of SP4206. When we divided the molecule into smaller and smaller fragments, this specificity was gradually lost, with the smallest fragments we tested (10 heavy atoms or fewer) being almost incapable of engaging the protein at specific binding sites or binding poses. This finding suggests that very small chemical fragments individually may not be capable of specific interaction with a protein. The distribution of the small fragments collectively on the protein surface in our simulations, however, yielded subtle yet discernable indications of the cryptic binding site’s location. To systematically identify cryptic binding sites using such small fragments, one could potentially develop an iterative method based on a set of spontaneous binding simulations that start with a limited set of small, basic chemical fragments, then combine them into progressively larger fragments, and ultimately into full-sized small molecules that hold the potential for selective binding to the target protein. This approach resembles the process of fragment-based lead discovery using X-ray crystallography,^46^ but with simulation instead of X-ray crystallography serving as the main platform.

By revealing atomistic details of the IL2-SP4206 binding process in high temporal resolution, our simulations shed new light on the process of small-molecule binding to a cryptic site. It has long been debated whether the chemical and geometric features of cryptic binding sites emerge as a result of interactions with a ligand (the induced fit model^47^) or, instead, whether these features emerge independently of a ligand, with the ligand only stabilizing them upon binding (the conformational selection model^48^). Our simulations suggest that the binding of SP4206 to IL2 may be best thought of as a hybrid process: The initial formation of an ion pair between the small molecule and the host protein exemplifies the mechanism of conformational selection, while the rest of the binding process—especially the full development of the binding groove—is consistent with an induced fit model.

## Acknowledgments

The authors thank Albert Pan and Michael Eastwood for helpful discussions and a critical reading of the manuscript, and Rebecca Bish-Cornelissen and Berkman Frank for editorial assistance.

## Supporting Information

Methods of simulation and analysis; movies of small-molecule binding.

## Notes

### Competing Interest Statement

The authors have declared no competing interest.

