## Supporting Information for "How does a small molecule bind at a cryptic binding site?"

### Supporting Methods

#### *MD simulations*

All IL2 simulations (Simulations 1–29; see Table S1) were based on an X-ray structure of IL2 (PDB 1PY2). Co-crystallized small molecules and non-protein atoms were removed, and the disordered loop regions were modeled to make a complete protein structure. The software package Maestro[1] was used to model the missing loops and side-chain atoms.

The IL2 simulation systems were set up by placing the protein at the center of the simulation box and filling the empty space with water and ion molecules. A cube with sides of 58–60 Å was used, resulting in system sizes of ~23,000 atoms. Na and Cl ions were added to maintain physiological salinity (150 mM) and to obtain a neutral total charge for the system. Most simulations contained a single copy of one of the small molecules (equivalent to ~8.0-mM concentration), with the exception of two simulations, as indicated in the text, in which we included three copies of SP4206 (equivalent to ~24.0-mM concentration) per simulation system with the intent of increasing the probability of binding. In the simulations of chemical-fragment binding, one copy of each fragment (equivalent to ~8.0-mM concentration) was included in the simulation systems shown in Figure 5A (fragments S and T), 5B (fragments L, M, and N), and

5D (18 fragments). Two copies of each fragment (equivalent to ~16.0-mM concentration) were included in the system shown in Figure 5C (fragments L, P, R, Q, and N). Each copy in the system translates into a ~9-mM concentration. In simulations of IL2 and fragments L, P, R, Q, and N, a higher concentration was chosen due to the weaker interactions of the smaller fragments with the protein.

The 18 fragments taken from ref. 2 and used in the simulations are amphetamine sulfate (molecular weight (MW) 135), hydroxyamphetamine hydrobromide (MW 151), benzoic acid (MW 122), salicylamide (MW 137), hydroquinone (MW 122), dopamine hydrochloride (MW 153), sulfanilamide (MW 172), xylose (MW 150), levorphanol tartrate (MW 257), prednisolone (MW 360), etilefrine (MW 181), ceftizoxime (MW 383), sodium oxybate (MW 104), cefetamet (MW 397), dextrose (MW 180), medazepam (MW 271), cyclizine lactate (MW 266), methamphetamine hydrochloride (MW 149).

The simulation systems of compound 43a (4-(4-Benzyl-4-methoxypiperidin-1-yl)-**N**-(4-(2-methyl-1-(phenylthio)propan-2-ylamino)-3-nitrophenylsulfonyl)benzamide) binding to Bcl-xL (Simulations 30–35; see Table S1) were set up and run similarly to the IL2 systems. Each simulation system contained one or three copies of the small molecule, initially randomly positioned away from the protein. The cubic simulation systems each contain 30,000 or fewer atoms.

In all the simulations, the systems were parameterized using the Amber99SB force field with corrections for Leu, Ile, Asp, and Asn[3] (which builds upon other modifications[4,5] to Amber99[6]) for the protein; TIP3P for water;[7] and GAFF with AM1-BCC for the small molecules.[8] We used the Anton specialized hardware[9] to perform all the simulations. The simulation methods and parameters are described in detail in ref. 10. Equilibrium MD simulations were performed on

Anton in the NVT ensemble at 310 K using the Nosé-Hoover thermostat[11] with a relaxation time of 1.0 ps. All bond lengths to hydrogen atoms were constrained using an implementation[12] of M-SHAKE.[13] Long-range electrostatic interactions were computed by the Gaussian split Ewald method.[14] A reversible reference system propagator algorithm (r-RESPA) integrator[15] was used to compute the bond terms, van der Waals and short-range electrostatic interactions, and long-range electrostatic interactions at different time intervals. A 2-fs time step was used for the bond terms and short-range interactions, and a 6-fs time step was used for the long-range interactions. For the Bcl-xL system, a 2-fs time step was used for bond terms and short-range interactions.

#### *FEP calculation*

In each spontaneous small-molecule binding simulation, when a small molecule remained at one location on the protein for more than 5 μs, it was deemed a potential binding event. A representative conformation of the conformations of the small molecule at the location was then obtained using a clustering algorithm,[16] and this conformation was considered to be the pose for this potential binding event. An absolute binding FEP calculation was then used to calculate the binding free energy of the small molecule to IL2 based on this binding pose. Depending on the small molecule, each FEP calculation involved a total simulation time of 5 μs, divided into 31–36 simulation windows, and was performed as described previously.[17,18,19]

#### *Other calculations*

The GBSA energy of binding shown in Figures 1E and S2 was calculated using Amber8.[20] The interaction energy, $[E_{\text{complex}} - E_{\text{protein}} - E_{\text{ligand}}] / 2$, for each snapshot was calculated using igb = 2 and cut = 300.  The occupancy maps of small molecules shown in Figure 5 were constructed using the volmap tool in VMD[21] using all heavy atoms, with the protein conformation aligned by the backbone αC atoms.  The volume of the binding groove was calculated using the MDPocket software.[22]

### Supporting Table, Figures, and Movies

**Table S1. Simulation details**

| | Sim # | Length (µs) | Protein | Small molecule(s) (S.M.) | # copies of S.M. | Specific Binding | Binding time range (µs) |
|---|---|---|---|---|---|---|---|
| SP4206 | 1 | 31.0 | IL2 | SP4206 | 3 | yes | 0.25— |
| | 2 | 21.0 | IL2 | SP4206 | 3 | no | |
| | 3 | 18.0 | IL2 | SP4206 | 1 | yes | 0.1— |
| | 4 | 25.0 | IL2 | SP4206 | 1 | no | |
| | 5 | 22.0 | IL2 | SP4206 | 1 | no | |
| | 6 | 17.0 | IL2 | SP4206 | 1 | no | |
| | 7 | 10.0 | IL2 | SP4206 | 1 | no | |
| | 8 | 10.0 | IL2 | SP4206 | 1 | yes | 1.05— |
| | 9 | 10.0 | IL2 | SP4206 | 1 | no | |
| | 10 | 10.0 | IL2 | SP4206 | 1 | yes | 0.43— |
| | 11 | 10.0 | IL2 | SP4206 | 1 | no | |
| | 12 | 10.0 | IL2 | SP4206 | 1 | yes | |
| | 13 | 10.0 | IL2 | SP4206 | 1 | yes | 0.62— |
| | 14 | 10.0 | IL2 | SP4206 | 1 | no | |
| | 15 | 10.0 | IL2 | SP4206 | 1 | yes | 8.0— |
| | 16 | 10.0 | IL2 | SP4206 | 1 | no | |
| SP4206 analogs | 17 | 20.0 | IL2 | SP4206-1 | 2 | yes | 16— |
| | 18 | 20.0 | IL2 | SP4206-1 | 2 | yes | 0.1— |
| | 19 | 20.0 | IL2 | SP4206-2 | 2 | yes | 1— |
| | 20 | 20.0 | IL2 | SP4206-2 | 2 | yes | 0.6— |
| | 21 | 20.0 | IL2 | SP4206-3 | 2 | no | |
| | 22 | 20.0 | IL2 | SP4206-3 | 2 | yes | 15— |
| | 23 | 20.0 | IL2 | SP4206-3 | 2 | no | |
| | 24 | 20.0 | IL2 | SP4206-3 | 2 | yes | 6.1— |
| fragments | 25 | 24.0 | IL2 | Fragments S, T | 2 | no | |
| | 26 | 53.9 | IL2 | Fragments S, T | 1 | yes | T(23.6—48.4) |
| | 27 | 37.0 | IL2 | Fragments N, M, L | 1 | yes | M(0.7—24.1), |
| | 28 | 16.2 | IL2 | Fragments N, Q, R, P, L | 2 | no | |
| | 29 | 16.5 | IL2 | Set of 18 fragments | 1 | no | |
| Compound 43a | 30 | 30.9 | Bcl-xL | Compound 43a | 1 | no | |
| | 31 | 36.0 | Bcl-xL | Compound 43a | 1 | no | |
| | 32 | 13.0 | Bcl-xL | Compound 43a | 1 | no | |
| | 32 | 38.0 | Bcl-xL | Compound 43a | 3 | yes | 31— |
| | 34 | 18.9 | Bcl-xL | Compound 43a | 3 | no | |
| | 35 | 11.7 | Bcl-xL | Compound 43a | 3 | yes | 3.1— |

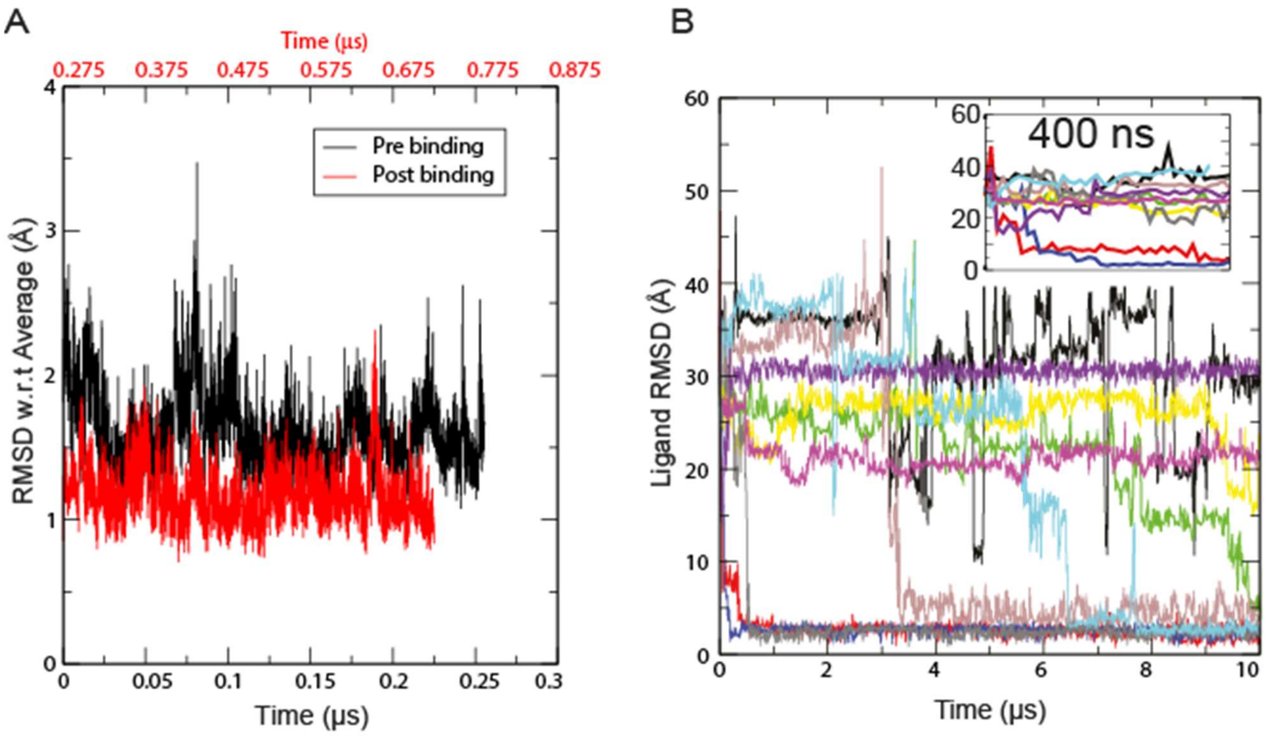


**Figure S1. Conformational fluctuation with respect to the average.** (A) This figure is a supplement to Figure 1D. The RMSD of the residues forming the binding groove (residues 61–65, 54, 55, 58, 59, 82, 85, 88, 92, 131) with respect to the average conformation of the period (pre-binding: 0–0.25 μs; post binding: 0.275–0.5 μs) is shown. As shown, the conformational fluctuation was reduced after SP4206 binding. (B) The time series of SP4206 RMSD with respect to the crystal binding pose in 10 additional simulations in which SP4206 was placed in a random initial position away from the protein. As shown, four simulations arrived at the native binding pose and two arrived at poses similar to the native. The inset is a magnification of the first 400 ns of the simulation.

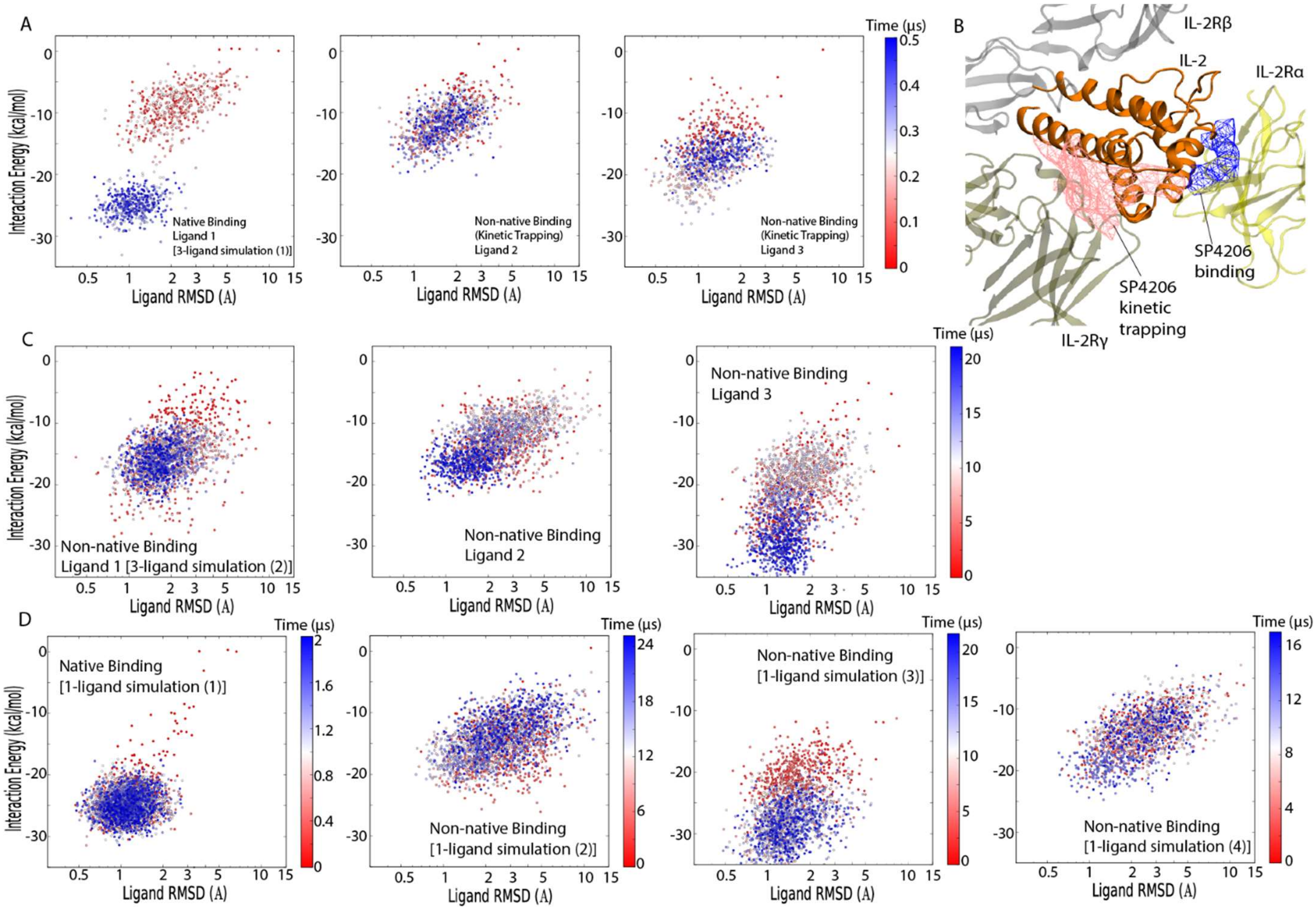


**Figure S2. Interaction energy of SP4206 with IL2 in simulations.** The estimated ligand-protein interaction energy (calculated with the GBSA method) and ligand conformational fluctuation were analyzed for six simulations. (A) A three-ligand simulation, in which one of the three ligands reached the native binding pose at 0.25 μs (Ligand 1), mapped to a 2D space of energy and ligand conformational fluctuation as measured by RMSD with respect to the conformation of the previous time step (*x*-axis). (B) The receptor domains (IL2Rα, β, and γ) bound with IL2 are shown with meshes indicating the high-occupancy regions for SP4206. (C) Another three-ligand simulation, in which none of the ligands reached the native pose. (D) Four one-ligand simulations. The ligand reached the native binding pose quickly (at 0.1 μs) in the first simulation, and as a result, the pre-binding dissociated state was under-sampled. In the other three simulations, the ligand did not reach the native pose.

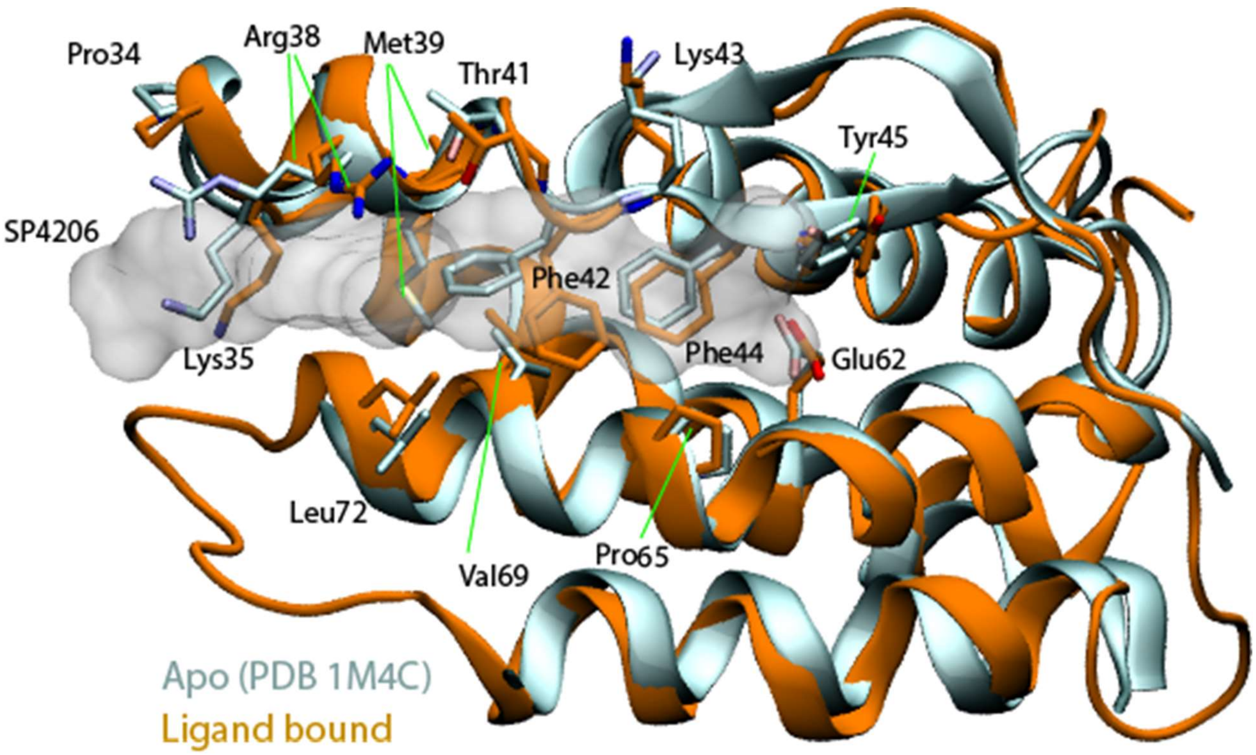


**Figure S3. Comparison of unbound and ligand-bound conformations of IL2.** The backbone ribbon of each of the two conformations is rendered, along with the residues in contact with the bound ligand, which are rendered as a transparent surface. The primary difference between the two conformations involves a side-chain rearrangement of the residues that form the binding site, particularly Arg38, Met39, and Phe42.

**Movie S1. Simulation of SP4206 binding to IL2.** A visualization of the simulation of SP4206 binding to IL2 detailed in Figure 1. IL2 is rendered with gray van der Waals spheres and SP4206 is rendered with orange licorice. The spontaneous emergence of the binding groove without SP4206 in the vicinity is highlighted. At the end of the movie, the simulation pose of SP4206 is superimposed on the crystallographic pose.

**Movie S2. Simulation of two SP4206 fragments and IL2.** A visualization of the simulation of two fragments (S and T) derived from SP4206 interacting with IL2 (detailed in Figure 5A).

**Movie S3. Simulation of three SP4206 fragments and IL2.** A visualization of the simulation of three fragments (L, M, and N) derived from SP4206 interacting with IL2 (detailed in Figure 5B).

**Movie S4. Simulation of five SP4206 fragments and IL2.** A visualization of the simulation of five fragments (L, P, R, Q, and N) derived from SP4206 interacting with IL2 (detailed in Figure 5C).

**Movie S5. Simulation of 18 chemical fragments and IL2.** A visualization of the simulation of 18 chemical fragments interacting with IL2 (detailed in Figure 5D).
